# (-)-Epigallocatechin-3-gallate inhibition of Epstein-Barr virus lytic replication involves latent membrane protein 1-mediated MAPK signaling pathways

**DOI:** 10.1101/383497

**Authors:** Hongde Li, Xiangjian Luo, Jianmin Hu, Sufang Liu, Namei Li, Min Tang, Xinxiang Weng, Wei Yi, Jinghe Gao, Ann M. Bode, Zigang Dong, Ya Cao

## Abstract

**Abstract:** EBV lytic replication has been shown to be important for carcinogenesis. Latent membrane protein 1 (LMP1) plays an important role in the viral latent infection and is abundantly expressed after EBV entry into the lytic cycle. However, the biological significance of LMP1 continuous expression in EBV lytic cycle is still not completely understood. We found that LMP1 promotes EBV reactivation by activating the downstream MAPK signaling in both AGS-EBV and B95.8 cells. In AGS-EBV cells, LMP1 induces EBV the initiation of the EBV lytic cycle in a p53 dependent manner. Activation of c-Jun by LMP1 through JNKs appears to be involved in EBV reactivation in p53 mutant B95.8 cells. We also demonstrated that EGCG, an anti-EBV agent, inhibits LMP1 expression and the activation of the downstream MAPK signaling pathways, followed by downregulation of EBV lytic protein expression level. Together, this study provides the first evidence that LMP1 promotes EBV reactivation via activation of the MAPK signaling pathways. Our findings further demonstrate that the mechanisms underlying EGCG inhibition of the EBV lytic replication involve the suppression of LMP1-mediated MAPK signaling pathways.

**Summary statement:** This study definitely confirms the role of LMP1 in EBV reactivation and further explores the mechanism by which EGCG inhibits EBV lytic replication.

## Introduction

Epstein-Barr virus (EBV) is the first human tumor virus identified and known to be linked to several types of human malignancies, including gastric carcinoma (GC), Hodgkin’s lymphoma (HL), Burkitt’s lymphoma (BL), and nasopharyngeal carcinoma (NPC)(Souza et al., 2005). In the host, EBV exhibits two alternative modes of infection, latent and lytic. In the latent infection, EBV only expresses a limited number of gene products such as latent membrane protein 1 (LMP1), LMP2A/2B, EBV nuclear antigens (EBNAs), and EBV-encoded small RNAs (EBERs). Studies indicated that the latent infection is a major cause of EBV-associated malignancies(Fiorina et al., 2014; Ko, 2015). LMP1 is a classic EBV-encoded oncoprotein and plays a critical role in *in vitro* B cell transformation. As mimics of CD40, LMP1 functions as a constitutively activated tumor necrosis factor receptor (TNFR) to activate multiple signaling pathways, including NF-κB(Ma et al., 2011b), PKC(Yan et al., 2007), JAK/STAT(Wang et al., 2010), MAPK(Li et al., 2007), and PI3-K/Akt(Mori and Sairenji, 2006) that contribute to its oncogenic effects by affecting cellular proliferation and survival.

When stimulated by certain chemicals or biological reagents, the latently infected cells will enter into the lytic phase of EBV infection. During the lytic cycle, EBV briefly passes through three consecutive lytic stages, including the immediate early (IE), early (E), and late (L), and first transcribes the IE genes, *BZLF1* and *BRLF1*, which encode two transactivators, Zta and Rta, respectively. The both proteins turn on the entire lytic viral cascade of gene expression, including *BMRF1* and *BALF5*, which encode diffused early antigen (Ea-D) and DNA polymerase. Simultaneously, Zta also acts as a replication factor for EBV genomic DNA by binding the lytic origin of replication, *oriLyt.* After that, the late lytic genes express the viral structural proteins, followed by the viral genome encapsidation and production of mature virions. The transition from the viral latent to lytic cycle is called EBV reactivation, which is essential for virus production and transmission from host to host.

Ahsan *et al.* reported that LMP1 plays a critical role in virus production and transmission(Ahsan et al., 2005). In this study, they provide the first evidence that LMP1 has a function in lytic reactivation. Based on the knowledge of mechanisms underlying EBV reactivation, LMP1 appears to facilitate EBV entry into the lytic cycle in a variety of ways. For instance, LMP1 leads to severe cellular stresses, which induce EBV lytic cycle initiation(Li et al., 2016), such as hypoxia(Benders et al., 2009), oxidative stress(Papadopoulou et al., 2014), and inflammation(Hannigan et al., 2011). In addition, LMP1-induced S-phase-like microenvironment is essential for lytic viral replication(Sueur et al., 2014). Thus, it is reasonable to speculate that LMP1 plays an important role in EBV reactivation.

LMP1 has been shown to enhance transcriptional activity and stability of p53 by inducing phosphorylation of p53 at Ser15, Ser392, Ser20, and Thr81 through MAPKs in NPC cells(Li et al., 2007). In an earlier study, Zhang *et al.* showed that p53 inhibits the ability of Zta to disrupt EBV latency(Zhang et al., 1994). However, some recent studies indicate that p53 is a prerequisite for EBV reactivation by facilitating the expression of Zta(Chang et al., 2008; Chua et al., 2012). Although it looks like controversial, these observations suggest the involvement of p53 in the regulation of EBV lytic cycle initiation.

Recently, EBV lytic replication has been shown to play an important role in carcinogenesis of several cancers. Clinical and epidemiological studies have revealed that individuals with elevated plasma EBV DNA load and antibody titers against the lytic viral capsid antigen (VCA) and early antigen (EA) have a high risk of EBV-induced malignancies(Jia and Qin, 2012; Liu et al., 2012). Also, recurrent EBV reactivation causes tumorigenesis by inducing oncogenic cytokines and enhancing genomic instability(Fang et al., 2009). Therefore, blocking EBV lytic replication is valuable for prevention and treatment of EBV-associated malignancies and helps to improve clinical outcome(Funch et al., 2005; Hong et al., 2005). EGCG, the major active polyphenol in green tea, is an inhibitor of EBV lytic cascade(Chang et al., 2003). Our previous study demonstrated that EGCG inhibits EBV spontaneous lytic replication through inactivation of MEK/ERK1/2 and PI3-K/Akt signaling(Liu et al., 2013).

In this study, we have investigated the contribution of LMP1 to EBV reactivation in EBV-positive gastric carcinoma and B-cell lymphoma cell lines and demonstrated that the MAPK signaling pathways, including ERKs, p38, and JNKs, are required for EBV reactivation by LMP1. In p53 wide-type AGS-EBV cells, LMP1 induces EBV reactivation in a p53-dependent manner. Activation of c-Jun by LMP1 through JNKs is involved in reactivation of EBV in p53 mutant B95.8 cells. We have also demonstrated that EGCG inhibition of EBV lytic replication is involved in the repression of the LMP1-mediated MAPK signaling pathways.

## Results

### Detection of EBV-encoded products in cell lines and biopsies

To investigate the possible role of LMP1 in the EBV lytic cycle, we used the AGS-EBV gastric carcinoma cell line containing recombinant EBV which is permissive for EBV lytic replication. We also used the B95.8 cells, a well-established EBV lytically-infected cell line. As shown, LMP1 and the lytic proteins, Zta and Ea-D, were expressed in both AGS-EBV and B95.8 cells (Figure 1A). By using a quantitative real-time PCR, the EBV genome copies were detected at significant levels in both AGS-EBV cells (5.34×10^6^ copies/μl) and B95.8 cells (6.27×10^6^ copies/μl) (Figure 1B).

**Figure 1.**
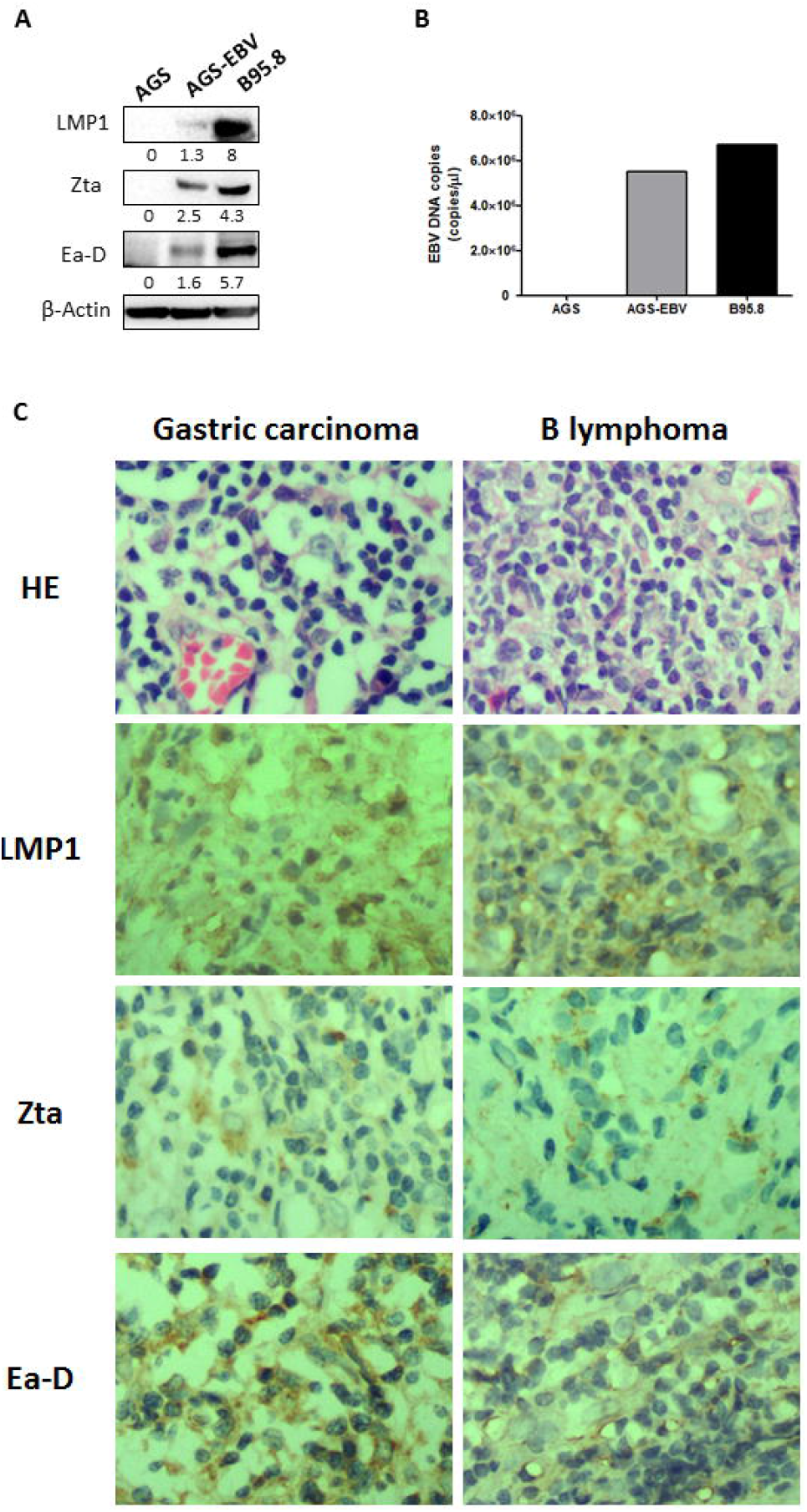
Detection of EBV-encoded proteins and EBV DNA in cell lines or/and biopsies of gastric carcinoma and B lymphoma. (A) Equal amounts of lysates were prepared from AGS, AGS-EBV, and B95.8 cells and western blot analysis was performed to determine the expression of LMP1, Zta, and Ea-D. AGS-EBV cell line was a negative control. (B) EBV genomic copies were isolated and determined by using a quantitative real-time PCR (RT-qPCR) assay with specific primers and a TaqMan probe targeting the BamHI W region of the EBV genome. (C) Paraffin sections were screened for the expression of EBV LMP1, Zta, and Ea-D with the specific antibodies. Positive signal implies that EBV LMP1, Zta, and Ea-D expressions occur in tumor cells as shown in different cases. Magnification, ×400.

Detecting EBERs by in situ hybridization has been regarded as a standard of EBV-positive in EBV-associated tumors. Immunohistochemistry to detect LMP1, Zta, and Ea-D was performed on formalin-fixed, paraffin-embedded tissue from EBER positive gastric carcinoma and B lymphoma. Positive signal implies that LMP1 expression occurs in tumor cells as shown by nuclear and/or cytoplasm localization, and expressions of Zta and Ea-D occur as shown by nuclear localization in different cases. We observed LMP1-, BZLF1-, and BMRF1-positive cells within the gastric carcinoma and B lymphoma biopsies. We detected the expressions of LMP1, Zta, and Ea-D in all (100%) of the 10 patients with EBER-positive gastric carcinoma and in all (100%) of 14 patients with EBER-positive B lymphoma (Figure 1C).

These results suggest that EBV spontaneous lytic replication exists in both cell lines and biopsies of EBV-positive gastric carcinoma and B lymphoma. Moreover, EBV-encoded LMP1 is also detected in the both biopsies.

### LMP1 promotes EBV reactivation in AGS-EBV and B95.8 cells

Then, we tested whether LMP1 could induce EBV reactivation. The cells were transfected with a plasmid containing pSG5-LMP1 or a control pSG5 vector. We found that Zta and Ea-D were markedly upregulated and EBV genome copies were higher compared to controls (Figure 2A). Next, knockdown of LMP1 by DZ1, a designed DNAzyme which specifically targets transcripts of LMP1, significantly reduced expression of Zta and Ea-D and decreased the numbers of EBV genome copy (Figure 2B).

**Figure 2.**
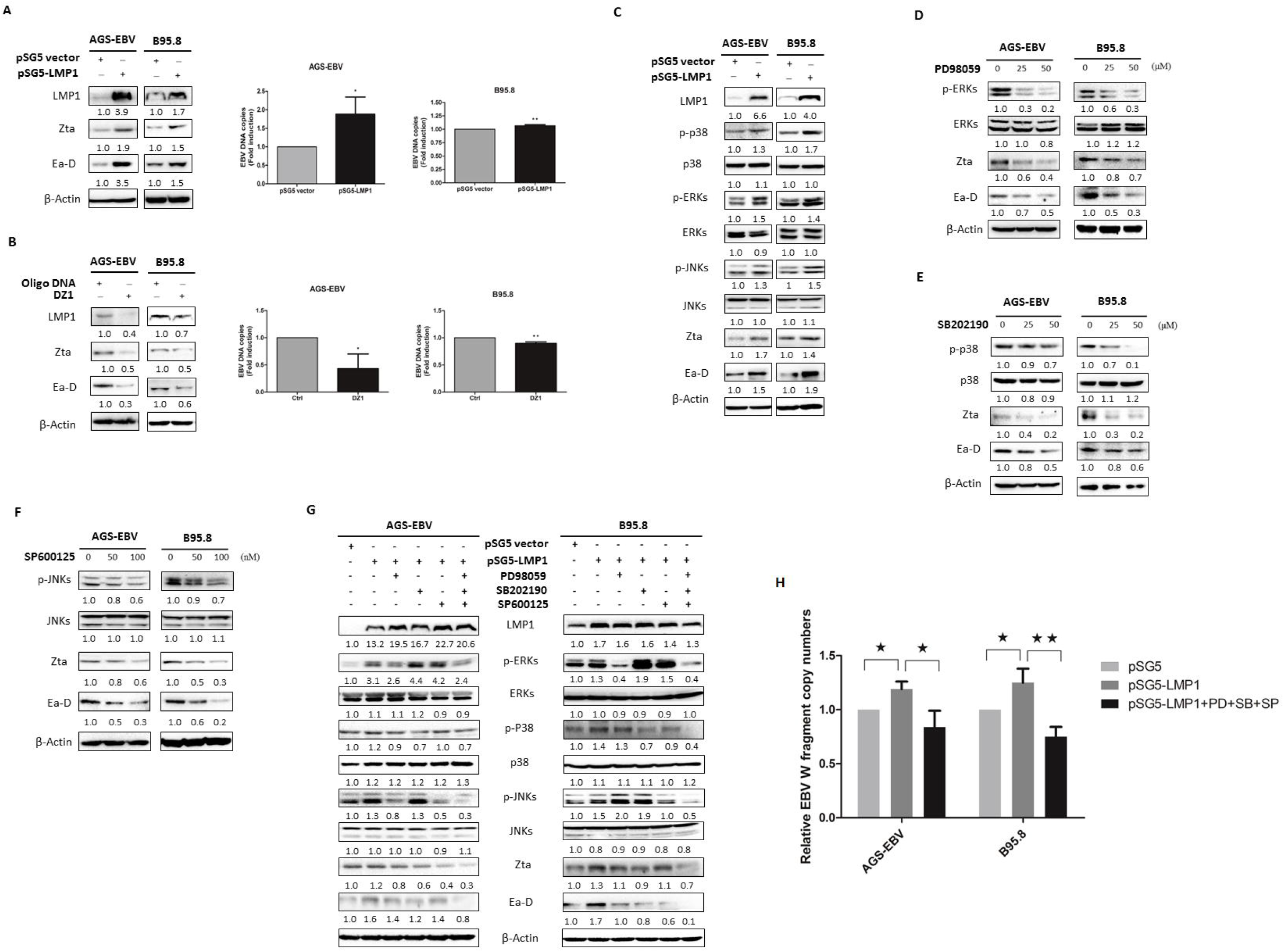
LMP1 promotes EBV reactivation via activating the downstream MAPKs. (A) AGS-EBV and B95.8 cells were transfected with a plasmid containing *pSG5-LMP1* or a control *pSG5* vector. Expression of LMP1, Zta, and Ea-D was detected. β-Actin was used as an internal control. EBV genome copies were determined by RT-qPCR. For each experiment, BamHI-W fragment levels from untreated cells were used as a control and set to 1. (B) AGS-EBV and B95.8 cells were transfected with DZ1 or random oligo DNA. The levels of LMP1, Zta, and Ea-D and numbers of EBV genome copies were detected. Each value represents the mean (± SD) of a representative of 3 independent experiments. Statistical significance: *p < 0.05, **p < 0.01 versus negative control. (C) AGS-EBV and B95.8 cells were transfected with a plasmid containing *pSG5-LMP1* or a control *pSG5* vector. Western blot analysis for LMP1, total and phosphorylated ERKs, p38, and JNKs, and EBV lytic proteins Zta and Ea-D were performed. (D, E, F) AGS-EBV and B95.8 cells were pretreated with MAPKs specific inhibitors, including PD98059 (0, 25, and 50 μM), SB202190 (0, 25, and 50 μM), and Sp600125 (0, 50, and 100 nM) for 24. Equal amounts of cell lysates were prepared and western blot analysis was performed to determine the levels of total and phosphorylated ERKs, p38, and JNKs and EBV lytic proteins. AGS-EBV and B95.8 cells were transfected with the plasmid expressing LMP1 for 4 h prior to incubation with MAPK specific inhibitors, including PD98059, SB202190, or/and SP600125, and then these cells were continuously cultured for 48 h. (G,H) The levels of total and phosphorylated MAPKs and lytic proteins and EBV genomic copy numbers were analyzed using western blot analysis and a quantitative real-time PCR (RT-qPCR) assay, respectively. Statistical significance: *p < 0.05, **p < 0.01 versus negative control. β-Actin was used as an internal control for A-H.

### MAPK signaling pathways are required for LMP1-induced EBV reactivation

Mitogen-activated protein kinases (MAPKs), including extracellular signal regulated kinases (ERKs), p38, and c-Jun NH2-terminal kinases (JNKs), are strongly activated in response to variety of different cellular stimuli (Goswami et al., 2012; Satoh et al., 1999). We found that phosphorylation levels of ERKs, p38, and JNKs were increased followed by LMP1 induction in AGS-EBV and B95.8 cells (Figure 2C). To determine whether MAPK signaling pathways is involved in LMP1-induced EBV reactivation, we first treated the cells with MAPK specific inhibitors. The inhibitors included PD98059, which is a specific inhibitor of MEK1 that acts by suppressing activation of ERK1/2, SB202190, a specific inhibitor of p38 kinase, and SP600125, an inhibitor of JNKs that exhibits a 300-fold greater selectivity for JNKs as compared to ERKs or p38. Pretreatment with PD98059, SB202190, or SP600125 markedly attenuated the expression of Zta and Ea-D in both cell models (Figure 2D, E, and F). To further verify the requirement of MAPK signaling pathways for EBV lytic cycle initiation induced by LMP1, the cells were transfected with a plasmid containing *pSG5-LMP1* or *pSG5* control and simultaneously incubated with PD98059, SB202190, or/and Sp600125. A significant reduction in Zta and Ea-D expression was detected compared to untreated controls. When the three inhibitors were simultaneously added to cells transfected with *pSG5-LMP1*, the protein levels of Zta and Ea-D were lower compared to the levels in cells treated with each individual inhibitor and were similar to the negative control. As expected, the levels of EBV genomic copies were lower compared to the levels in cells transfected with pSG5-LMP1 (Figure 2G, H).

Together, these observations above suggest that LMP1 promotes EBV reactivation and the activation of the downstream MAPK signaling pathways is required for LMP1-induced EBV reactivation.

### LMP1 induces EBV reactivation in a p53-dependent manner in p53 wide-type AGS-EBV cells

P53 is an important downstream molecule of the MAPK signaling pathways. To examine the role of p53 in EBV lytic initiation, AGS-EBV and B95.8 cells were transfected with the plasmid expressing wide-type p53. We found that inducible p53 expression significantly increased the levels of Zta and Ea-D (Figure 3A). We further knocked down p53 by siRNA. The data showed that Zta and Ea-D were downregulated in AGS-EBV cells, but no significant changes were detected in B95.8 cells (Figure 3B). As p53 is mutant in B95.8 cell line (Farrell et al., 1991; Forte and Luftig, 2009; Li et al., 2008), and there is no hot spot mutation of p53 in AGS-EBV cell line (Supplementary Figure 1), suggesting that wild-type p53 may be critical for regulation of EBV reactivation. Moreover, inducible expression of p53 also increased numbers of EBV genomic copy (Figure 3C). Our results further showed that knockdown of p53 by siRNA counteracted LMP1-induced expressions of both Zta and Ea-D and EBV DNA replication in AGS-EBV cells (Figure 3D, E). Next, we tested the effect of p53 on the activity of *BZLF1* promoter (Zp) using a Zp/luciferase reporter system in combination with transfection of the plasmid expressing p53 in p53-null H1299 cells. As shown, p53 activated Zp in a dose dependent manner (Figure 3F). A ChIP assay showed that p53 could bind to the positions -221 to +12 of Zp in AGS-EBV cells (Figure 3G). Recently, studies have shown that p53 interacts with the universal transcription factor Sp1 to activate Zp(Chua et al., 2012) and is also known to interact with members of the SMAD family of transcription factors to enhance transcription from their downstream target genes(Overstreet et al., 2014). By using a co-immunoprecipitation (Co-IP) assay, our results showed that the interactions between p53 and Sp1 or SMAD2 were significantly enhanced after LMP1 induction (Figure 3H). Thus, LMP1 may improve the ability of p53 to recruit Sp1 and SMAD2 on sits of Zp, inducing Zta expression and thus EBV reactivation.

**Figure 3.**
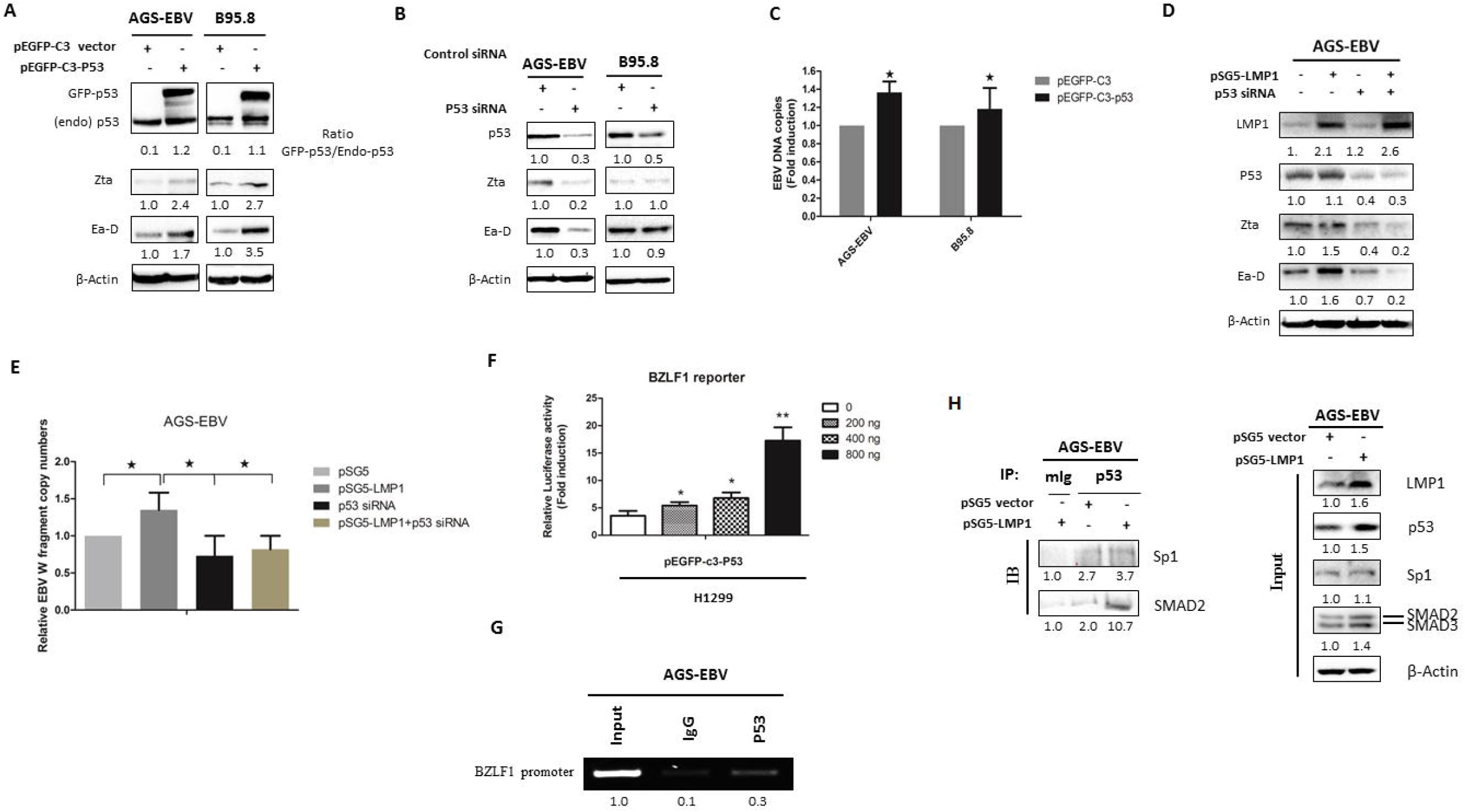
P53 promotes EBV reactivation by binding to and activating *BZLF1* promoter (Zp) as well as through interaction with Sp1/Smad2. (A) AGS-EBV and B95.8 cells were transfected with a plasmid containing *pSG5-LMP1* or a control *pSG5* vector for 48 h. Equal amounts of cell lysates from the cells were prepared and western blot analysis was performed to determine expression of EGFP-p53, endogenous p53, and EBV lytic proteins. (B) The cells were transfected with siRNA against *p53* or control siRNA for 48 h. Western blot analysis was performed to determine expression of EGFP-p53, endogenous p53, and EBV lytic proteins. β-actin was used as an internal control. (C) AGS-EBV and B95.8 cells were transfected with a plasmid containing *pSG5-LMP1* or a control *pSG5* vector for 48 h. EBV genome copies were analyzed using RT-qPCR with specific primers designed to amplify the EBV W fragment. For each experiment, BamHI-W fragment levels from untreated cells were used as a control and set to 1. Each value represents the mean (± SD) of a representative of 3 independent experiments. Statistical significance: *p < 0.05, **p < 0.01, *** *p* < 0.001 versus negative control. (D, E) AGS-EBV cells were co-transfected with plasmid expressing LMP1 and p53 siRNA for 48 h. *pSG5* empty vector and control siRNA were added as controls. Cell lysates were harvested and subjected to western blot analysis for LMP1, p53, and lytic proteins, EBV genomic copy numbers were analyzed using a quantitative real-time PCR (RT-qPCR) assay. Statistical significance: *p < 0.05, **p < 0.01 versus negative control. (F) H1299 cells were co-transfected with the Zp reporter and plasmid expressing p53 or empty vector for 48 h. The transfected cells were then harvested for determining the effect of p53 on activity of the *BZLF1* promoter (Zp) luciferase activity assay. The relative activity was normalized to the value of the internal control *pRL-SV40* plasmid activity. Each value represents the mean (± SD) of a representative of 3 independent experiments. Statistical significance:*p < 0.05, **p < 0.01 versus negative control. (G) AGS-EBV cells were subjected to analyze the interaction of p53 with Zp using ChIP assay. The regions of positions -221 to +12 of Zp were amplified by PCR. The relative intensities of anti-p53-precipitated Zp DNA amounts were provided after being normalized with those of their input controls. P21 promoter was used as a positive control. (H) AGS-EBV cells were transfected with plasmid expressing LMP1 or empty vector for 48 h. The cells were harvested for co-immunoprecipitation (co-IP) assay using anti-p53 antibody to analyze the interactions of p53 with Sp1 and SMAD2/3. The resulting immunocomplexes were recovered and analyzed by western blot analysis. β-Actin was used as an internal control for A-H.

The results suggest that wide-type p53 induces Zta expression by binding to and activate Zp and that LMP1 enhances the ability of p53 to activate Zp via directly binding to Zp as well as through cooperation with Sp1, promoting Zta-mediated EBV reactivation in AGS-EBV cells.

### Phosphorylation of p53 and c-Jun mediated by MAPKs are involved in LMP1-induced EBV reactivation in AGS-EBV and B95.8 cells, respectively

In a previous study, our group demonstrated that LMP1 induces phosphorylation of p53 through MAPKs in NPC cells (Li et al., 2007). Phosphorylation of p53 at Ser15 by ERKs results in p53 for transactivation, phosphorylation of p53 at Ser392 by p38 regulates its oligomerization and affinity for sequence-specific DNA, and phosphorylation of p53 at Ser20 by JNKs is crucial for p53 stabilization. Here, inducible expression of LMP1 increased the levels of total p53, phosphorylated MAPKs, and phosphorylated p53 at Ser15, Ser20, and Ser392 in AGS-EBV cells (Figure 4A). After treatment with PD98059 and SB202190, phosphorylation of p53 at Ser15 and Ser392 was reduced as result of the inactivation of ERKs and p38, respectively, and no significant changes in p53 expression were detected, followed by the downregulation of Zta and Ea-D expression (Figure 4B, C). P53 expression and phosphorylation of p53 at Ser20 were both decreased after inactivation of JNKs by SP600125, followed by the downregulation of Zta and Ea-D expression (Figure 4D). The results suggest that LMP1 regulates the transcriptional function and stability of p53 through MAPKs in AGS-EBV cells, implying that LMP1 enhances the ability of p53 to promote Zta-mediated EBV reactivation.

**Figure 4.**
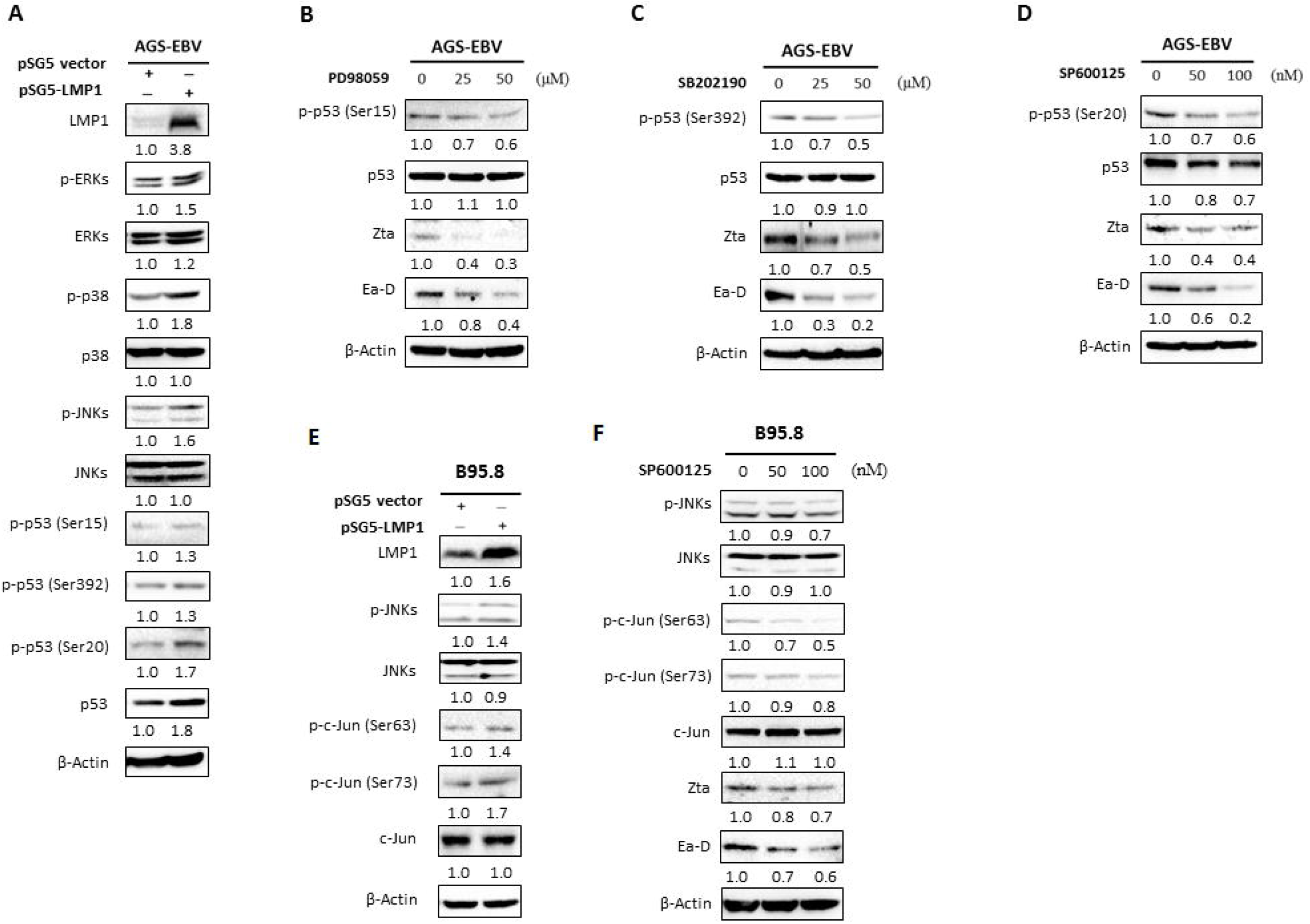
Phosphorylation of p53 and c-Jun by LMP1 through MAPKs plays an important role in EBV reactivation in p53 wide-type AGS-EBV cells and p53 mutant B95.8 cells, respectively. (A) AGS-EBV cells were transfected with a plasmid containing *pSG5-LMP1* or a control *pSG5* vector. for 48 h. Then the cells were harvested for western blot analysis for LMP1, total MAPKs and p53, and phosphorylated MAPKs and p53. (B, C, and D) AGS-EBV cells were pretreated with PD98059, SB202190, or Sp600125 for 24 h at the concentrations indicated. Western blot analysis for EBV lytic proteins and phosphorylated p53 was performed. (E) B95.8 cells were transfected with a plasmid containing *pSG5-LMP1* or a control *pSG5* vector. Cell lysates were harvested and western blot analysis was performed to analyze LMP1, total c-Jun and JNKs, and phosphorylated c-Jun and JNKs. (F) B95.8 cells were pretreated with different concentrations of Sp600125 (0, 50, and 100 nM) for 24 h. Western blot analysis for c-Jun, phosphorylated c-Jun, and EBV lytic proteins were performed. β-Actin was used as an internal control.

MAPKs is also shown to modify and activate many transcription factors of Zp, such as c-Jun, ATFs, and CREB (Kenney and Mertz, 2014; Li et al., 2016). C-Jun is a component of the transcription factor activator protein-1 (AP-1) complex(Feng et al., 2007), and its transcriptional activity is regulated by phosphorylation at Ser63 and Ser73 through JNKs (Morton et al., 2003). In B95.8 cells, we found that the levels of phosphorylated JNKs and phosphorylated c-Jun at Ser63 and Ser73 were increased after LMP1 induction (Figure 4E). Furthermore, inactivation of JNKs mediated by SP600125 inhibited phosphorylation of c-Jun at Ser63 and Ser73, followed by the decreased expression of Zta and Ea-D (Figure 4F). The evidences indicate that activation of c-Jun by LMP1 through JNKs might be involved in reactivation of EBV in B95.8 cell line which has p53 mutation.

### The mechanism by which LMP1 induces EBV reactivation in AGS-EBV and B95.8 cells

LMP1, as mimics of CD40, recruits TRAFs(Korchnak et al., 2009) to constitutively activate MAPKs, including ERKs, p38, and JNKs. In AGS-EBV cells, LMP1 induces phosphorylation of p53 through MAPKs to enhance its transcriptional function and improve its enrichment. P53 activates *BZLF1* promoter (Zp) by directly binding to Zp as well as through cooperation with Sp1 on Zp, leading to Zta expression. Then, Zta trigger EBV reactivation. In p53 mutant B95.8 cells, activation of c-Jun by JNKs is involved in LMP1-induced reactivation of EBV and this mechanism should be considered to participate in EBV reactivation by LMP1. Thus, MAPK specific inhibitors, including PD98059, SB202190, and SP600125, effectively inhibit LMP1-induced EBV (Figure 5).

**Figure 5.**
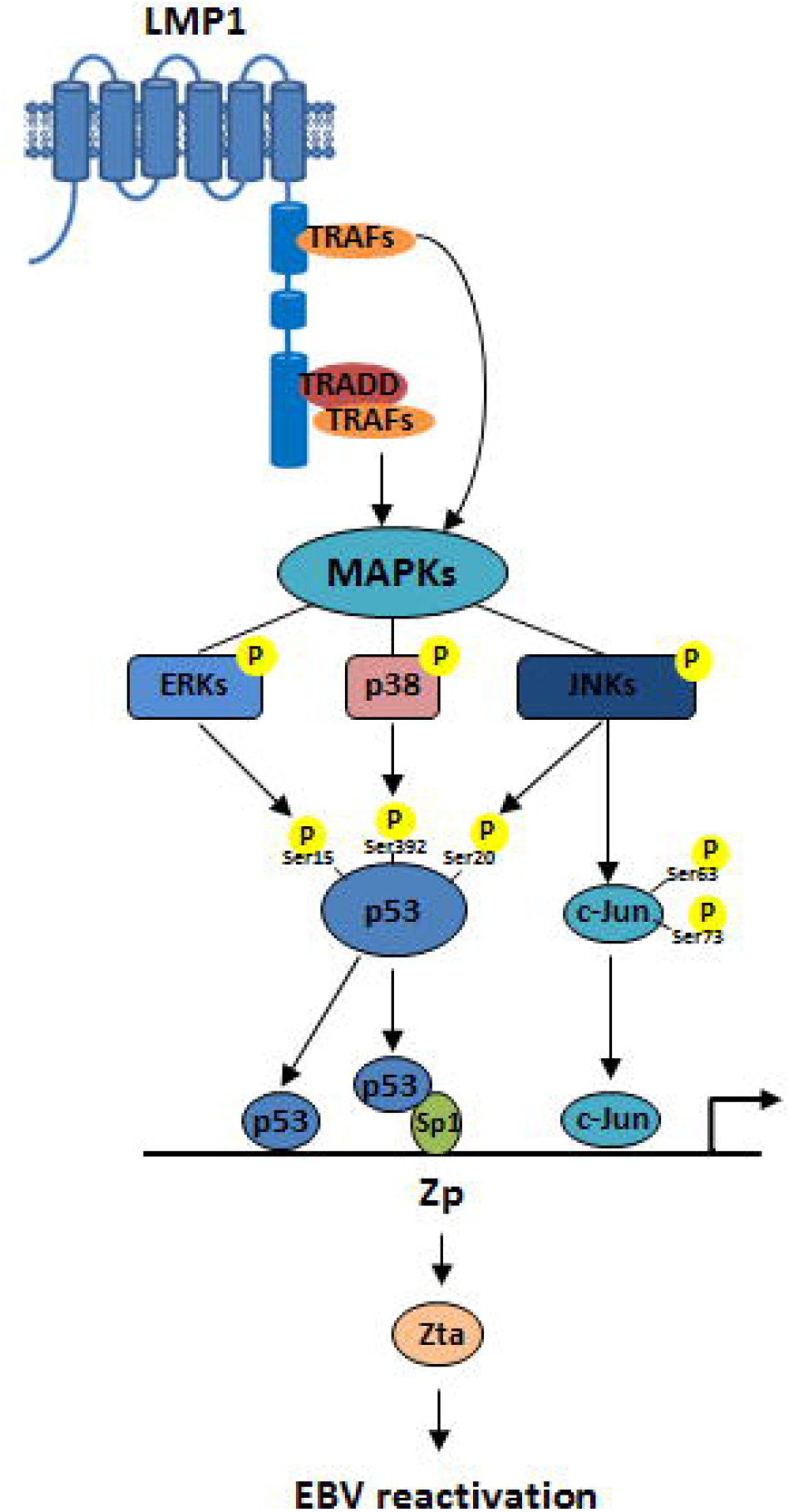
Schematic illustrating the role of LMP1 in inducing EBV reactivation in AGS-EBV and B95.8 cells.

### EGCG inhibits EBV spontaneous lytic replication by suppressing LMP1-midiated MAPK signaling pathways

Our recent study demonstrated that EGCG inhibition of EBV spontaneous lytic infection involves repression of PI3-K/Akt and MEK/ERKs signaling pathways in EBV-positive NPC cells(Liu et al., 2013). To further explore the mechanisms by which EGCG inhibits EBV reactivation, AGS-EBV and B95.8 cells were incubated with EGCG. Results showed that EGCG treatment significantly downregulated Zta and Ea-D expression (Figure 6A). Consistent with the results, RT-qPCR assay showed that transcripts of lytic genes, *BZLF1* and *BMRF1*, were substantially attenuated by EGCG (Figure 6B). In addition, we detected a reduction in the number of EBV genome copies after EGCG treatment (Figure 6C). Flow cytometry analysis revealed that after treatment with EGCG, the percentage of cells expressing Zta decreased from 16.24 to 10.93% in AGS-EBV cells and from 17.51 to 5.79% in B95.8 cells. Additionally, the percentage of cells expressing Ea-D decreased from 34.80 to 15.18% in AGS-EBV cells and from 27.67 to 14.77% in B95.8 cells (Figure 6D). These data confirm the inhibitory effects of EGCG on EBV spontaneous lytic replication in the both cell lines.

**Figure 6.**
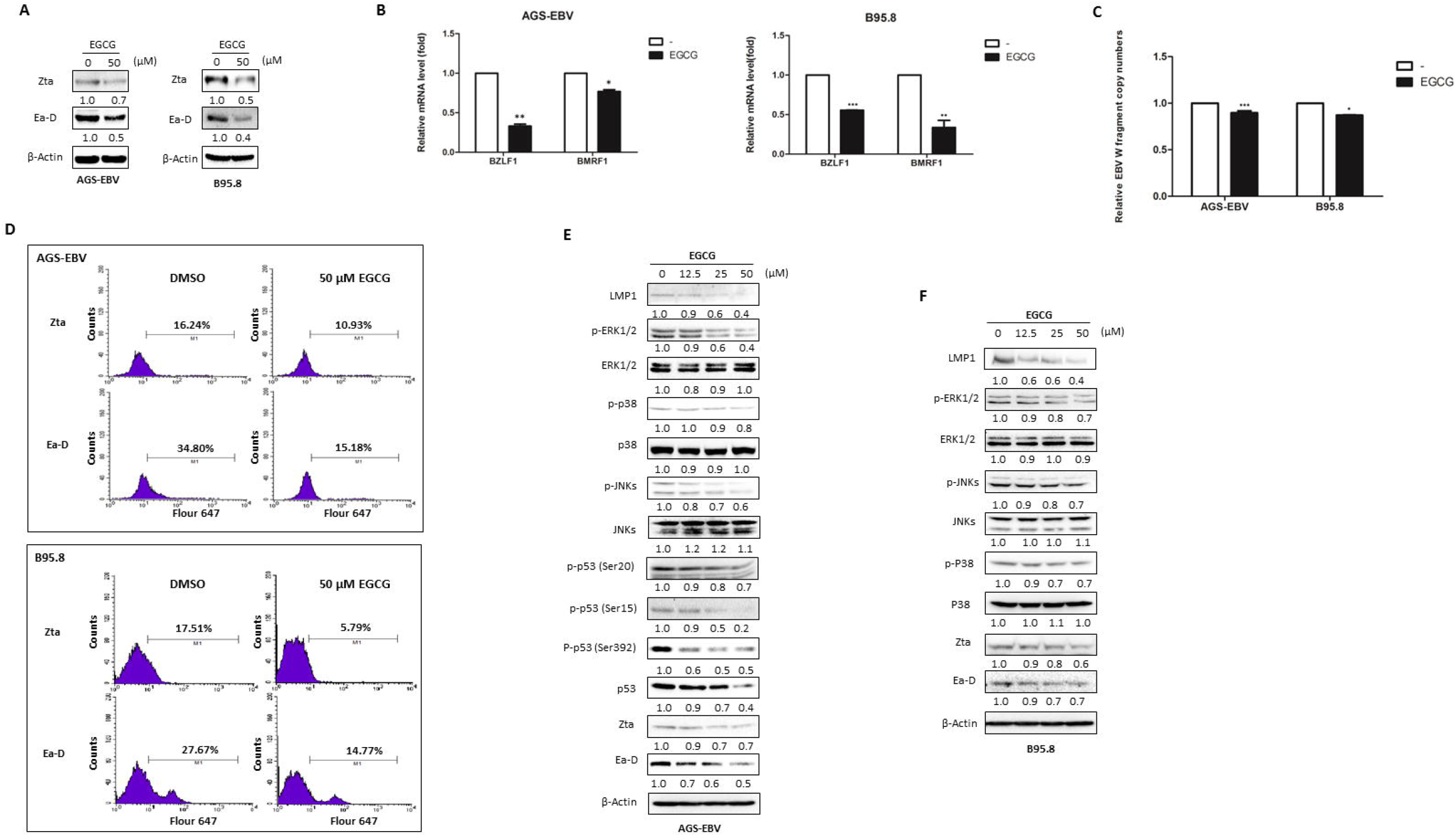
EGCG inhibition of EBV lytic replication involves the suppression of LMP1-mediated MAPK signaling pathways in AGS-EBV and B95.8 cells. AGS-EBV and B95.8 cells were starved in 0.1% FBS HAM’S/F-12 or RPMI medium 1640 for 24 h and then treated with 50 μM EGCG or vehicle (DMSO) for 24h. (A) Equal amounts of cell lysates were prepared and western blot analysis was performed to determine the expression of Zta and Ea-D. β-Actin was used as an internal control. (B) An RT-qPCR assay was performed to determine the effects of EGCG on the mRNA levels of *BZLF1* and *BMRF1* genes. (C) The changes in EBV genome copies were analyzed using RT-qPCR with specific primers designed to amplify the EBV W fragment. For each experiment, BamHI-W fragment levels from untreated cells were used as a control and set to 1. Each value represents the mean (± SD) of a representative of 3 independent experiments. Statistical significance: *p < 0.05, **p < 0.01, *** *p* < 0.001 versus negative control. (D) Flow cytometry was performed to analyze the effects of EGCG on the numbers of Zta- or Ea-D-expressing cells. Incubation with vehicle but no primary antibody followed by incubation with the secondary antibody was used as a negative control. AGS-EBV and B95.8 cells were starved in 0.1% FBS HAM’S/F-12 or RPMI medium 1640 for 24 h and then treated with different concentrations of EGCG as indicated or vehicle (DMSO) for 24h. (E and F) Cell lysates were harvested and analyzed for LMP1, total and phosphorylated MAPKs, and lytic proteins using western blot analysis. Effects of EGCG on the levels of total and phosphorylated p53 were also determined in AGS-EBV cells. β-Actin was used as an internal control.

Furthermore, EGCG was shown to reduce LMP1 expression and phosphorylated ERKs, p38, and JNKs in a dose-dependent manner, followed by downregulation of Zta and Ea-D expression (Figure 6E, F). In AGS-EBV cells, total p53 and phosphorylated p53 were significantly downregulated by EGCG in a dose-dependent manner (Figure 6E). Moreover, a co-IP assay showed that EGCG inhibited the ability of p53 to bind Sp1 (Supplementary Figure 3). While in B95.8 cells, EGCG treatment decreased the levels of total c-Jun and phosphorylated c-Jun in a dose-dependent manner (Figure 6F). Thus, this observation suggests that EGCG could inhibit EBV spontaneous lytic infection by repressing LMP1-mediated MAPKs/p53 signal axis in AGS-EBV cells and JNK MAPK/c-Jun axis in B95.8 cells.

## Discussion

Over the past 20 years, EBV-attributable malignancies have increased by 14.6%(Khan and Hashim, 2014). As known, EBV has been associated with gastric carcinoma (GC), Hodgkin’s lymphoma (HL), Burkitt’s lymphoma (BL), and nasopharyngeal carcinoma (NPC). Among them, GC, especially gastric adenocarcinomas, was first categorized by EBV-positivity (9%)(Cancer Genome Atlas Research, 2014). Recently, it has been widely reported that EBV lytic replication contributes to carcinogenesis, and the current knowledge points to the fact that EBV spontaneous lytic replication takes place in variousmethod after normalization EBV-associated tumors. Our results further indicated that the EBV spontaneous lytic infection exists in both gastric adenocarcinomas and B lymphoma. The results from immunohistochemistry showed co-expression of Zta and Ea-D in EBER-positive gastric adenocarcinomas and B lymphoma biopsies, and the relative expression of the both proteins is consistent with the cell level (Figure 1).

LMP1 is known to intensify EBV latent infection and also abundantly expressed during the viral lytic cycle. A recent study of recombinant EBV has been demonstrated that LMP1 plays a critical role in virus production and release(Ahsan et al., 2005). LMP1 was expressed either in cell lines and biopsies of the both tumors (Figure 1A and 1C). Therefore, LMP1 may be involved in initiation of the EBV lytic cycle. In this study, we showed that LMP1 contributes to EBV reactivation in both AGS-EBV and B95.8 cells (Figure 2). Furthermore, our results showed that the MAPK specific inhibitors synergistically counteracted LMP1 induction of lytic protein expression (Figure 2G), indicating that the MAPK signaling pathways are required for LMP1-induced EBV reactivation.

P53 is deemed to be involved in regulation of EBV reactivation, our results confirmed that p53 contributes to EBV reactivation, and wide-type p53 is critical for the effect (Figure 3A and 3B). And we further found that p53 binds to and activates *BZLF1* promoter (Figure 3G). Besides, we also demonstrated that p53 binds to and activate another IE gene promoter, *BRLF1* promoter (positions -442 to -2) (Supplementary Figure 2). The evidence suggests that p53 may induce expression of Zta and Rta by binding to and activating Zp/Rp, amplifying its lytic-inducing effects on EBV lytic cycle initiation.

Li *et al.* showed that phosphorylation of p53 at Ser15 by ERKs results in p53 for transactivation, phosphorylation of p53 at Ser392 by p38 regulates its oligomerization and affinity for sequence-specific DNA, and phosphorylation of p53 at Ser20 by JNKs is known to be crucial for p53 stabilization(Li et al., 2007). In AGS-EBV cells, LMP1 was shown to induce phosphorylation of p53 at Ser15, Ser392, and Ser20 and increase p53 protein level through MAPKs (Figure 4A, B, C, and D), which suggest that LMP1 might regulate the ability of p53 to bind to and activate Zp. P53 is shown to promote the transcription from Zp via interaction with Sp1(Chua et al., 2012). In addition, p53 cooperates with SMADs to transactivate downstream target genes(Cordenonsi et al., 2003). And we detected the enhanced interactions between p53 and transcription factors Sp1 or SMAD2 after LMP1 induction (Figure 3H), suggesting that phosphorylation of p53 by LMP1 also promotes p53 enrichment and thus improves the ability of p53 to recruit Sp1 and SMAD2 on sites of Zp. However, it needs to be investigated whether the cooperation of p53 with SMAD2 contributes to EBV reactivation. Furthermore, we found that knockdown of p53 by siRNA significantly counteracts LMP1-induced expression of lytic proteins (Figure 3D). These results demonstrate that LMP1 contributes to EBV lytic cycle initiation in a p53-dependent manner in AGS-EBV cells. In p53 mutant B95.8 cells, activation of c-Jun by LMP1 through JNKs could be involved in EBV reactivation (Figure 4E and 4F). We speculate that LMP1 may also promote EBV reactivation by inducing activation of other downstream transcription factors through MAPKs.

Altogether, as mimics of CD40, LMP1 recruits TRAFs(Korchnak et al., 2009) to constitutively activate the MAPK signaling pathways, including ERKs, p38, and JNKs. In AGS-EBV cells, LMP1 induces phosphorylation of p53 at Ser15, Ser20, and Ser392 to enhance the ability of p53 to activate Zp by directly binding to Zp as well as through interaction with Sp1, promoting Zta-mediated EBV reactivation. In p53 mutant B95.8 cells, LMP1-induced activation of c-Jun through JNKs is involved in reactivation of EBV and this mechanism should be considered to participate in EBV reactivation by LMP1. Thus, blocking the MAPK signaling pathways by specific inhibitors effectively inhibits EBV lytic replication (Figure 5).

The EBV lytic cycle has been shown to be important for carcinogenesis, and some lytic proteins are known to have carcinogenic properties. For example, Zta induces the production of several oncogenic and inflammatory cytokines (Katsumura et al., 2012; Ma et al., 2011a; Ma et al., 2012). BGLF4 protein kinase (EBV-PK) and BGLF5 nuclease (EBV DNase), have been reported to promote genomic instability and enhance progression of cancer cells(Chang et al., 2012; Wu et al., 2010). A recent study showed that BALF3 effectively induces genomic instability and mediates NPC relapse(Chiu et al., 2014).Thus, blocking EBV lytic replication has been developed as an effective preventive and therapeutic strategy for EBV-associated malignancies.

In our previous study, we have demonstrated that EGCG blocks EBV spontaneous lytic replication by suppressing the activation of MEK/ERK1/2 and PI3-K/Akt signaling in NPC cells(Liu et al., 2013). Here, we further found that EGCG inhibition of EBV lytic replication involves blocking LMP1-mediated MAPK signaling pathways (Figure 6). As known, EGCG interacts with and binds numerous proteins to prevent carcinogenesis. Proteins that can directly bind with EGCG include the plasma proteins, such as fibronectin, Fas, laminin and the 67 kDa laminin receptor, vimentin ZAP-70, Fyn, insulin-line growth factor-I receptor (IGF-IR), and glucose-regulated protein 78 (GRP-78) (Bode and Dong, 2009). Recently, we have demonstrated that EGCG binds LMP1 from lysates in both AGS-EBV and B95.8 cells or purified from *E.coli* expression system by Ni-His tag (unpublished data), implying that EGCG may also affect LMP1-induced EBV reactivation by directly binding LMP1.

In summary, this study first demonstrated that LMP1 promotes EBV reactivation by activating the MAPK signaling pathways. The observation provides a novel insight into LMP1 link the EBV lytic infection. We further demonstrated that the mechanisms by which EGCG inhibits EBV lytic replication involve the repression of LMP1-mediated MAPK signaling pathways, supporting the development of EGCG as an anticancer and chemoprevention agent for EBV-associated malignancies in the future.

## Materials and methods

### Cell culture

The human gastric adenocarcinoma cell line AGS (CRL-1739™) and AGS-EBV containing recombinant EBV were cultured in HAM’S/F-12 medium supplemented with 10% fetal calf serum. B95.8 (VR-1492™), an EBV-positive B-lymphoma cell line, was cultured in RPMI1640 medium supplemented with 10% new born calf serum. Non-small cell lung carcinoma cell line H1299 (CRL-5803™) was cultured in RPMI1640 medium supplemented with 10% fetal calf serum. All cells were incubated in a humidified atmosphere of 5% CO_2_ at 37°C. AGS-EBV cells were provided by Prof. Wenhai Feng (the State Key Laboratory of Agrobiotechnology, China Agricultural University) and were authenticated by analyzing the DNA sequencing data of genes deleted or mutated (see supplementary material). AGS, B95.8, and H1299 were obtained from the ATCC and maintained as per the recommendations.

### Chemicals

EGCG (E4143), PD98059 (P215), and SB202190 (S7067) were purchased from Sigma Chemical (St. Louis, MO). SP600125 (HY-12041) was purchased from Medchem Express (Princeton. NJ, USA), and Protease Inhibitor Cocktail (B14002) was purchased from Selleck Chemicals (HOU, USA). All drugs were dissolved in DMSO and used at the indicated concentrations.

### RT-qPCR

Total RNA were isolated from cultured cells with TRIzol^®^ total RNA extraction reagent (Invitrogen, Carlsbad, CA) according to the manufacturer’s instructions, followed immediately by reverse transcription (RT), which was performed on 2μg of total RNA using the RevertAid First Strand cDNA Synthesis Kit (Thermo, USA) in a final volume of 20 μl. Quantitative PCR (qPCR) was performed using Fast Start Universal SYBR Green Master (ROX) and 2μl (10 × dilution) of RT reaction products (Roche, Basel, CH) with the primers of *BZLF1* and *BMRF1*. The primers are detailed in Supplementary table 1. The following PCR program was used: 95°C for 10 min; 95°C for 15 s, 60°C for 1 min (plate read), 40 cycles; melt curve 60-95°C, increments of 0.5°C, for 15s. Relative gene expression was calculated using the 2^−ΔΔct^ method after normalization to the reference gene β-actin.

### Western blot analysis

Cell lysates (50 μg) were separated by SDS–PAGE gel and transferred to polyvinylidene difluoride (PVDF) membranes. The membranes were incubated with blocking buffer for 1 h and then with appropriate dilutions of each primary antibody in blocking buffer overnight at 4°C. After washing the membrane 3 times (5 min each) with tris-buffered saline with Tween^®^ 20 (TBST), the membrane was incubated with the recommended dilution of the conjugated secondary antibody in blocking buffer at room temperature for 1 h. After incubation, the membrane was washed 3 times (5 min each) with TBST and then the signal was developed with the Super Signal^®^ West Femto Stable Maximum Sensitive Substract kit (Thermo, USA) following the kit manufacturer’s recommendations. The signal was detected by Molecular Imager^®^ ChemiDoc™ XRS+ (BIO-RAD, California, USA).

### Immunohistochemistry (IHC)

Ten paraffin-embedded EBER-positive gastric carcinoma tissues and 14 paraffin-embedded EBER-positive B lymphoma tissues were obtained from the Department of Pathology at Hunan Tumor Hospital and Xiangya Hospital, Changsha, China. Immunohistochemistry was performed on 4.0 μm sections from formalin-fixed, paraffin-embedded tissues. The sections were deparaffinized and rehydrated. Antigen retrieval was achieved with diluted ethylenediaminetetraacetic acid (50:1) by microwaving for 10 min. Endogenous peroxidase activity was quenched with a 3% hydrogen peroxide block for 15 min at room temperature. The sections were treated with goat serum for 10 min and incubated at 4°C overnight with the primary antibody. The secondary antibody, biotinylated horse anti-mouse IgG, was applied for 30min at 37°C, followed by 30 min incubation with Vectastain ABC Elite. The reaction was visualized by using diaminobenzidene chromogen (Dako) following the manufacturer’s instructions, and the slides were counterstained with Mayer’s hematoxylin. All slides were scored by two observers. The staining intensity and pattern were evaluated using a 0 to 3+ scale (0, completely negative; 1+, weak; 2+, intermediate; 3+, strong). A final score of ≥2+or greater was required for the case to be considered positive.

### Co-Immunoprecipitation (Co-IP) Assay

Immunoprecipitation was performed with AGS-EBV cells that were transfected with a plasmid containing the *pSG5* vector or *pSG5-LMP1*. After 72 h, cell lysates (500μg) prepared with IP lysis buffer were incubated with 20 μl protein A sepharose beads (Sigma, St. Louis, MO) for 2 h at 4 °C on a rotating rocker and centrifuged for 10 min at 2000 rpm to preclear. The recovered supernatant fraction was incubated with anti-p53 (sc-126 X, Santa Cruz, CA) in the presence of 1× protease inhibitors at 4°C overnight on a rotating rocker. Then 40 μl of protein A sepharose beads were added, and the incubation was continued for 2 h at 4°C on a rotating rocker. The immunocomplexes bound to sepharose beads were recovered by a brief centrifugation followed by 3 washes with cold 1× PBS. The harvested beads were resuspended in 20 μl of 5× SDS PAGE sample buffer and boiled for 5 min to release the bound proteins. A 50 μg cell lysate was used as an input control. The samples were analyzed by Western blotting.

### Chromatin immunoprecipitation (ChIP) assay

ChIP assays were performed essentially as described by Shi Y (Shi et al., 2012). Cells (5×10^6^) were cross-linked with 37% formaldehyde (1% final volume, Sigma). Cross-linking was stopped by the addition of 1/10 volume of 1.25 M glycine. Then the cells were harvested and disrupted using cell lysis buffer. The lysates containing chromatin were sonicated to obtain DNA fragments with an average length of <500 bp. After centrifugation, the protein concentration of the lysates was determined by the Bradford assay using the Pierce^™^ BCA Protein Assay Reagent (Thermo, USA). The protein-chromatin (300 μg) complexes were used for each immunoprecipitation. The antibody-protein complexes were captured with pre-blocked protein G dynabeads (Invitrogen, Carlsbad, CA). Immunoprecipitation DNA was analyzed by PCR for the quantification of DNA sequences and the recovered DNA was amplified by PCR. ChIP primers are detailed in Supplementary table 1.

### DNA extraction and quantification of EBV copy number

DNA was extracted from each sample (2 ×10^6^ cells) using the QIAamp^®^DNA Mini Kit (QIAGEN, Chatsworth, CA) according to the kit handbook. To quantify EBV copy numbers, a 10-fold dilution from 10^7^ to 10^4^ copies was first used to create the standard curve according to the instructions included with the EBV Polymerase PCR Fluorescence Quantitative Diagnostic Kit (DA, China). Then 2μl of each DNA sample were added to PCR reaction tubes, and PCR was performed as follows (ABI-7500, Applied Biosystems, USA): 93for 2 min, 93°C for 45 s, and 55°C for 1 min for 10 cycles; 93°C for 30 s and 55°C for 45 s (data collection) for 30 cycles. The EBV copy number of each sample can be calculated by the corresponding threshold (CT) cycle with the aid of the standard curve.

### Flow cytometer analysis

AGS-EBV and B95.8 cells (1~2×10^6^) were treated with EGCG (50 μM) for 24 h and then collected and washed with phosphate-buffered saline (PBS) to achieve a single cell suspension. This was followed by fixing with 4% paraformaldehyde for 30 min at 4°C. After washing with PBS, the fixed cells were permeabilized for 15 min with PBS containing 0.1% Triton X-100. The cells were then washed with PBS and blocked with 2% BSA and 0.1% Triton X-100 in PBS for 30 min at room temperature. The cells were washed with PBS and incubated with 1:100 diluted monoclonal Zta antibody (Abcam, Cambridge, UK) and monoclonal EA-D antibody(Abcam, Cambridge, UK) for 2 h at 4°C. Next, the cells were washed with PBS and incubated with 1:100 diluted Alexa Fluor^®^ 633-conjugated goat anti-mouse IgG (Life Technologies, Carlsbad, CA) for 1 h at room temperature. Incubation with PBS, without primary antibody followed by incubation with the secondary antibody, was used as a negative control. Finally, the cells were washed with PBS and resuspended in 1% paraformaldehyde for analysis using a FACSCalibur flow cytometer (BD Biosciences, USA).

### Transfection and luciferase assay

The cells were seeded into 24-well plates at the appropriate density and allowed to adhere overnight. When cells reached 70~90% confluence, various amounts of *pEGFP-C3-p53* were co-transfected respectively with *pZp-luc* or *pGL3-basic* and the *pRL-TK* internal control into cells using the Lipofectamine^¯^ 2000 reagent (Invitrogen, Carlsbad, CA) according to the manufacturer’s instructions. After transfection for 6 h, medium was replaced with fresh complete medium without antibiotics. The cells were harvested 48 h later and lysates were prepared.

### Statistical analysis

Data are expressed as mean values ± S.D. and differences between various groups were determined using the Student’s *t*-test. A value of *p*<0.05 was considered statistically significant.

## Acknowledgements

We thank Prof. Qiao Wu for giving p53 expression plasmid as a gift, Prof. Wenhai Feng for giving the AGS-EBV cells, and Prof. Chung S. Yang for valuable suggestions.

## Competing interests

No competing interests declared

## Funding

This work was supported by the NSFC/NIH pilot project (81161120410), the National Natural Science Foundation of China (81672705 and 81101474), the Innovation Foundation of Central South University (2013zzts072), the State Key Laboratory of Medicinal Chemical Biology (201611001), and the Collaborative Innovation Center for Chemistry and Molecular Medicine of Hunan province, China.

